# Single-cell atlas of a non-human primate reveals new pathogenic mechanisms of COVID-19

**DOI:** 10.1101/2020.04.10.022103

**Authors:** Lei Han, Xiaoyu Wei, Chuanyu Liu, Giacomo Volpe, Zhifeng Wang, Taotao Pan, Yue Yuan, Ying Lei, Yiwei Lai, Carl Ward, Yeya Yu, Mingyue Wang, Quan Shi, Tao Wu, Liang Wu, Ya Liu, Chunqing Wang, Yuanhang Zhang, Haixi Sun, Hao Yu, Zhenkun Zhuang, Tingting Tang, Yunting Huang, Haorong Lu, Liqin Xu, Jiangshan Xu, Mengnan Cheng, Yang Liu, Chi Wai Wong, Tao Tan, Weizhi Ji, Patrick H. Maxwell, Huanming Yang, Jian Wang, Shida Zhu, Shiping Liu, Xun Xu, Yong Hou, Miguel A. Esteban, Longqi Liu, South China Greater Bay Area-Single Cell Consortium (SC-GBA-C)

## Abstract

Stopping COVID-19 is a priority worldwide. Understanding which cell types are targeted by SARS-CoV-2 virus, whether interspecies differences exist, and how variations in cell state influence viral entry is fundamental for accelerating therapeutic and preventative approaches. In this endeavor, we profiled the transcriptome of nine tissues from a *Macaca fascicularis* monkey at single-cell resolution. The distribution of SARS-CoV-2 facilitators, ACE2 and TMRPSS2, in different cell subtypes showed substantial heterogeneity across lung, kidney, and liver. Through co-expression analysis, we identified immunomodulatory proteins such as IDO2 and ANPEP as potential SARS-CoV-2 targets responsible for immune cell exhaustion. Furthermore, single-cell chromatin accessibility analysis of the kidney unveiled a plausible link between IL6-mediated innate immune responses aiming to protect tissue and enhanced ACE2 expression that could promote viral entry. Our work constitutes a unique resource for understanding the physiology and pathophysiology of two phylogenetically close species, which might guide in the development of therapeutic approaches in humans.

**Bullet points:**

1. We generated a single-cell transcriptome atlas of 9 monkey tissues to study COVID-19.
2. *ACE2^+^TMPRSS2^+^* epithelial cells of lung, kidney and liver are targets for SARS-CoV-2.
3. *ACE2* correlation analysis shows *IDO2* and *ANPEP* as potential therapeutic opportunities.
4. We unveil a link between IL6, STAT transcription factors and boosted SARS-CoV-2 entry.

## INTRODUCTION

As the distance between humans and wild animal habitats diminishes due to uncontrolled human expansion, a series of zoonotic diseases with high mortality rates have emerged. For instance, the recent outbreak of Ebola in Africa, which killed over 5,000 people, was most likely spread from bats and primates to humans^1^. The current outbreak of coronavirus disease 2019 (COVID-19) caused by the coronavirus severe acute respiratory syndrome coronavirus 2 (SARS-CoV-2)^2^ is not the only example of coronaviruses that have recently passed from animals to humans. Coronaviruses are a family of RNA viruses that typically cause respiratory tract infections in humans, yet they are frequently in the reservoir of wild animals with no disease^3^. For example, the common cold is often (10-15%) caused by a coronavirus (e.g. HCoV-229E and HCoV-OC43)^4^. However, coronaviruses can also lead to severe and life-threatening diseases. In the early 2000s a coronavirus called SARS-CoV, believed to be passed from bats to humans in South East Asia, caused more than 700 deaths from around 8,000 confirmed cases worldwide^5^. Since 2012, another zoonotic coronavirus believed to have passed from camels to humans in the Middle East was designated as Middle East Respiratory Syndrome (MERS)^6^. To date, there have been over 2,500 confirmed cases of MERS with over 800 deaths. While SARS appears to have been eradicated, MERS cases are sporadic and human to human spread is limited^4^.

As of 21^st^ April 2020, COVID-19^2^ has become a global pandemic with more than ~2,500,000 confirmed cases and over 170,000 deaths. Due to its high infectivity rate and the high level of intensive care that many patients need, COVID-19 has overwhelmed national health services and destabilized the world. One important reason is that many people who are positive for the virus show mild symptoms^7, 8^, leading to unnoticed spread of the virus. The current worldwide emergency, possibility of continued expansion to less developed countries, risk of virus mutations and the perpetuation beyond this season has made it imperative to stop the trajectory of virus spreading. Developing drugs and preventative vaccines are ongoing but to warrant success it is necessary to have more knowledge about the disease mechanisms. So far, little is known except for the viral binding via angiotensin converting enzyme 2 (ACE2) and subsequent priming by type 2 transmembrane serine protease 2 (TMPRSS2) protease, which are shared mechanisms with SARS and MERS^9, 10^. To test experimental treatments, animal models close to humans are necessary due to sequence variation of ACE2 and changes in the proportions of cell subtypes in organs between species. For these reasons, it is essential to have a species close to human to study COVID-19. In this regard, monkey experiments have shown that infection with SARS-CoV-2 produces clinical manifestations similar to COVID-19 patients^11^. Another study demonstrated that infection with SARS-CoV-2 in monkeys is preventable by acquired immunity, answering one of the outstanding questions about the disease^12^.

Issues about the proportions of cell types within organs between species and their crosstalk can be addressed effectively through single-cell profiling technologies, in particular single-cell RNA-sequencing (RNA-seq) and single-cell assay for transposase accessible chromatin-sequencing (ATAC-seq). Yet, although human data are accumulating^13^, monkey data are still scarce. The comparison between human and monkey data will be crucial for advancing our knowledge of COVID-19. Here, we provide a high-resolution single-cell atlas of nine organs/tissues (lung, kidney, pancreas, brain, parotid, liver, thyroid, aorta artery, and blood) in monkey, encompassing 215,334 cells. By comparing the expression of SARS related targets in monkey and human, we have identified cell-to-cell similarities as expected. Crucially, we also discovered stark differences in *ACE2* expression between these two species, for example in the ciliated vs pulmonary alveolar type 2 cells of the lung and hepatocytes in liver. We also observed that *ACE2* is heterogeneous among different epithelial cell subtypes across these organs/tissues, suggesting that variations in cell state could influence viral entry. Supporting this, single-cell ATAC-seq of monkey kidney identified regulatory elements driven by signal transducer and activator of transcription factor 1 and 3 (STAT1 and 3) and interferon regulatory factor 1 (IRF) in the *ACE2* locus. This suggests that cytokines, particularly interleukin 6 (IL6), aiming to induce a tissue protective response can exacerbate the disease by aiding viral entry into target cells. Additionally, through correlation analysis with *ACE2* expression, we have identified several potential candidates involved in COVID-19 pathophysiology, such as Transmembrane protein 27 (TMEM27), Indoleamine 2,3-dioxygenase 2 (IDO2), DnaJ heat shock protein family (Hsp40) member C12 (DNAJC12) and Alanyl aminopeptidase N (ANPEP). These targets may offer therapeutic opportunities.

Taken together, our data constitute a unique resource which could aid the scientific community in the fight against SARS-CoV-2. From a wider perspective, this will also be instrumental for systematic comparative studies aimed at understanding physiological and pathophysiological differences between monkey and other species, in particular, human.

## RESULTS

### Cellular heterogeneity of nine non-human primate tissues assessed by single-cell RNA-seq

We profiled, at the single-cell level, the transcriptome of the model organism cynomolgus monkey *(Macaca fascicularis)*, as it is phylogenetically close to human and this could help advance our knowledge of human physiology and disease. As proof of principle, we decided to use our data to understand what cell types are mainly targeted by SARS-CoV-2 and how this could trigger the clinical features that have been lethal in a number of patients. For this study, we used a six-year-old female monkey in which we profiled nine different organs (**Fig. 1a**). These included lung, liver and kidney as the known affected organs by the closely related SARS-CoV infection^14^, and have been reported to have high *ACE2* expression in human^15^. Peripheral blood mononuclear cells (PBMC) were added because altered immune responses are thought to be detrimental in the disease^16^. Neocortex was chosen because of the clinical symptoms which involve loss of smell and taste suggesting that the central nervous system may be targeted^17^. The parotid gland was chosen on the basis that saliva is one of the main means of infection spread. Additionally, we selected aorta, thyroid and pancreas.

**Fig. 1.**
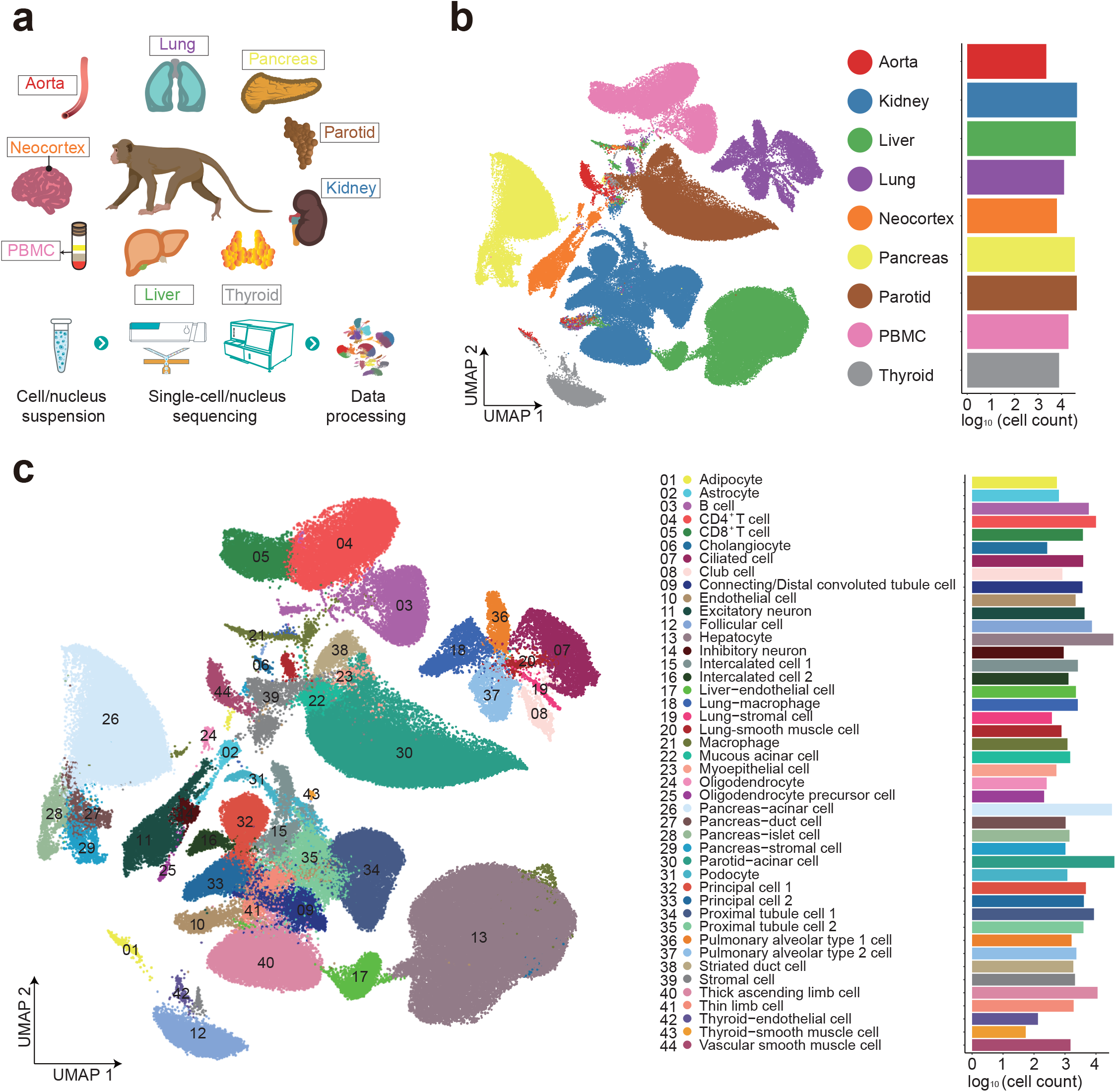
Construction of single-cell atlas across nine tissues of a *Macaca fascicularis* monkey. **a**, Schematic representation of selected monkey tissues used in this study and description of experimental pipeline for the single-cell sequencing. **b**, UMAP visualization of all single cells from the dataset colored by tissue/organ (left) and number of cells from each tissue passing quality control (right). **c**, UMAP visualization of each cell type colored according to 44 clusters in the first round of clustering. Cell type annotation is provided in the figure and is associated with a number indicative of every cluster. n = 215,334 individual nuclei/cells.

We employed a high-throughput platform recently developed in-house, DNBelab C4, which is a scalable and cost-effective approach for microfluidic droplet-based approach^18^. Except for PBMC sequencing, which was performed using cells in suspension, the sequencing for all the other organs was done using single-nucleus library preparations. Following euthanasia, the selected organs were extracted, single-nucleus/cell suspensions were obtained and then used for library preparation. A total of 40,226 liver, 45,286 kidney, 36,421 pancreas, 44,355 parotid gland, 12,822 lung, 7,877 thyroid, 6,361 neocortex, 2,260 aorta nuclei and 19,726 PBMCs passed quality control and were used for downstream analysis (**Extended Data Fig. 1a, b, Supplementary Table 1**).

In a global view of our single-cell dataset, each organ clustered separately, with the exception of a few cell types such as macrophages, adipocytes and endothelial cells, which were shared between different organs (**Fig. 1b**). We performed Uniform Manifold Approximation and Projection (UMAP) on the 215,334 cells and identified 44 major clusters by performing unbiased graph-based Louvain clustering (**Supplementary Table 1**). Some clusters were largely composed of cells belonging to a specific tissue, such as hepatocytes in cluster 13, pancreatic acinar cells in cluster 26 and parotid acinar cells in cluster 30 (**Fig. 1c, Extended Data Fig. 1c**). We next performed clustering and differential gene expression analysis to dissect the cellular composition of each individual organ. These analyses confirmed the typical patterns of cell heterogeneity for all the organs/tissues. When examining the lung tissue, we defined 10 major clusters with specific molecular markers, including ciliated cells, macrophages, cycling macrophages, smooth muscle cells, fibroblasts, pericytes, pulmonary alveolar (pneumocytes) type 1 and type 2, endothelial and club cells (**Extended Data Fig. 2a**). The kidney consisted of 11 clusters, those being podocytes, thick ascending limb cells, proximal tubule cells, intercalated cells 1 and 2, connecting tubule cells, distal convoluted tubule cells, stomal cells, thin limb cells, principal cells and endothelial cells (**Extended Data Fig. 2b**). Analysis of liver tissue revealed hepatocytes to be the larger cell population, while other clusters consisted of cholangiocytes, macrophages (Kupffer cells), natural killer-T (NK-T) cells, endothelial cells and hepatic stellate cells (**Extended Data Fig. 2c**). Inspection of PBMC clustering revealed large populations of B cells, CD4^+^, CD8^+^ naïve and CD8^+^ memory T cells, together with smaller populations of natural killer (NK) cells, dendritic cells, CD16^+^ and CD14^+^ monocytes (**Extended Data Fig. 2d**). Likewise, the neocortex contained excitatory neurons, astrocytes, microglia, parvalbumin (PVALB), somatostatin-expressing neurons (SST), synaptic vesicle glycoprotein-expressing cells (SV2C), vasoactive intestinal polypeptide-expressing neurons (VIP), oligodendrocytes and oligodendrocyte precursor cells (**Extended Data Fig. 2e**). Parotid gland instead was composed of a large cluster of serous acinar cells together with small clusters of macrophages, stromal cells, myoepithelial cells, striated duct cells, mucous acinar cells and intercalated duct cells (**Extended Data Fig. 2f**). Aorta cells could be further divided into adipocytes, endothelial cells, myofibroblasts and a large proportion of smooth muscle cells (**Extended Data Fig. 2g**). Our clustering also demonstrated that most of the thyroid gland is composed of follicular cells, with smaller populations of adipocytes, endothelial cells, stromal and smooth muscle cells (**Extended Data Fig. 2h**). Finally, our data showed the largest population of the pancreas to be acinar cells, while smaller clusters were comprised of stromal, ductal, and islet cells (alpha and beta), together with a population that could not be assigned to any known cell type (**Extended Data Fig. 2i**).

In conclusion, we have successfully profiled the transcriptome of nine organs at single-cell resolution in monkey, which could assist in the study of COVID-19.

### *ACE2* and *TMPRSS2* single-cell expression landscape in a non-human primate

Recent studies have reported that, similarly to SARS-CoV, the capacity of SARS-CoV-2 virus to infect host cells relies on viral spike (S) protein binding to ACE2 entry receptor^9, 10^, which is involved in the control of blood pressure through the renin-angiotensin system^19^. This phenomenon is primed by the multifunction serine protease TMPRSS2^20^. Accordingly, double positive (ACE2^+^/TMPRSS2^+^) cells have higher risk of infection by SARS-CoV-2. Although immunohistological studies have demonstrated localization of these two proteins in the respiratory tract^21^, it is unclear which cell subtypes express these genes and how homogenous the expression among a specific cell subtype is. Also, comprehensive information about other cell types and organs that express these two proteins and could be targeted by the virus in human or monkey is lacking.

We inspected our data to see how widespread and homogenous *ACE2* expression was in the monkey tissues. As expected, *ACE2* was detected in several lung clusters, mainly ciliated cells, club cells and pulmonary alveolar type 2 cells (**Fig. 2a, 2d, 3a upper panel**), whereas in the kidney, *ACE2* was primarily present in proximal tubule cells (**Fig. 2a, 2d, 3b upper panel**). The latter is consistent with reports describing that a significant number of COVID-19 patients display altered kidney function^15, 22^. Interestingly, *ACE2* expression was heterogenous among these cell subtypes in both lung and kidney. In the liver, *ACE2* was mostly expressed in cholangiocytes, with a smaller degree of expression also found in hepatocytes (**Fig. 2a, 2d, 3c upper panel**). Notably, the closely related SARS-CoV caused liver injury due to hepatitis in some patients^23^, suggesting that the liver may also be a direct target for SARS-CoV-2. A small proportion of *ACE2*^+^ was also observed in pancreatic islet cells (**Fig. 2a, 2d, Extended Data Fig. 3a**). In contrast, little or no expression was observed in thyroid, neocortex, parotid and PBMC (**Fig. 2a, 2d, Extended Data Fig. 3a**). Negligible *ACE2* expression in the neocortex suggests that other tissues may be affected by SARS-CoV-2 that cause loss of taste and smell, regarding the latter in particular the olfactory epithelium.

**Fig. 2.**
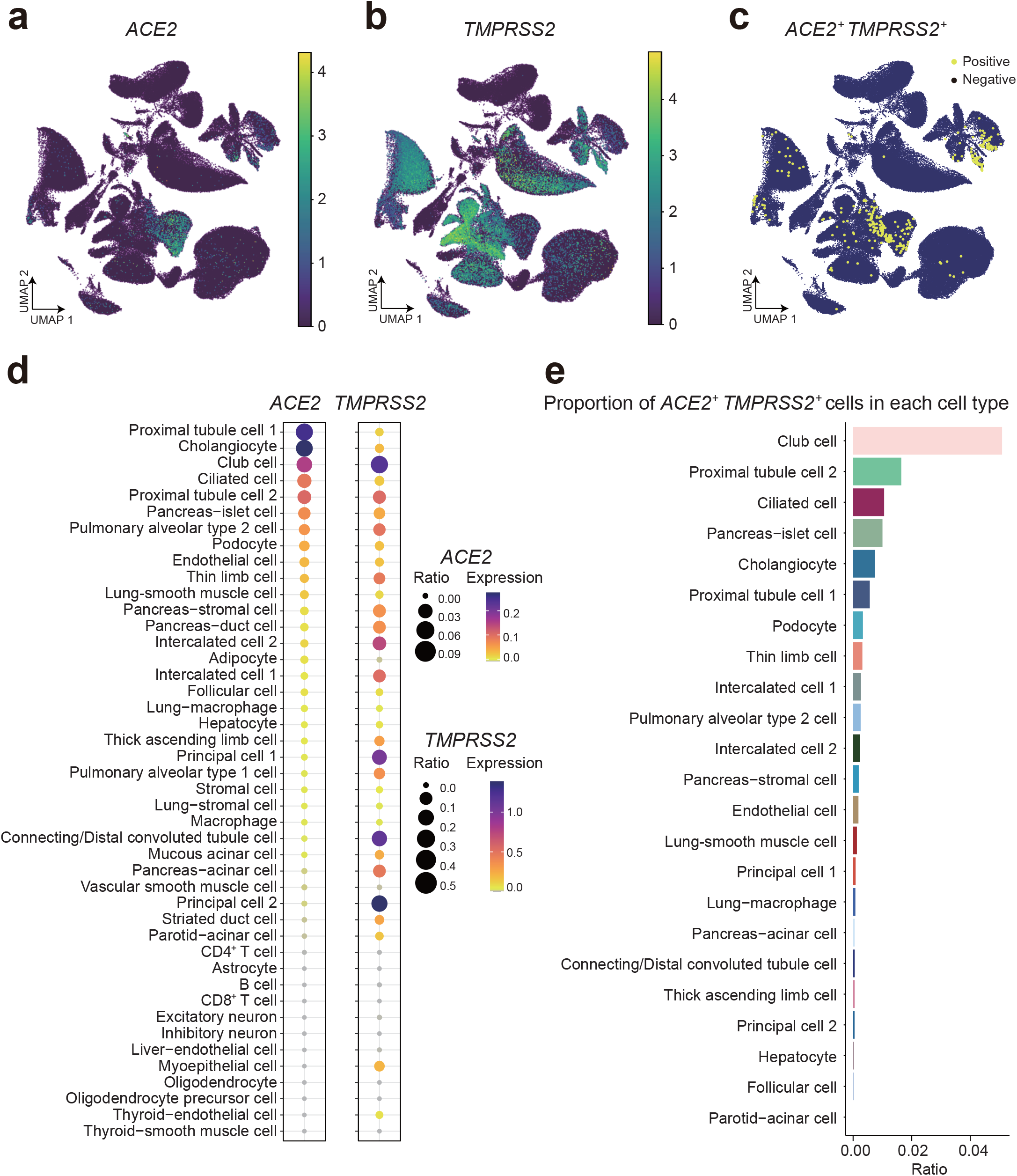
*ACE2* and *TMPRSS2* expression across 44 cell clusters in monkey. **a-b**, UMAP projection of *ACE2* (**a**) and *TMPRSS2* (**b**) expression in all single cells within our dataset. **c**, UMAP projection of *ACE2*^+^/*TMPRSS2*^+^ cells. **d**, Bubble plots showing the level of expression of *TMPRSS2* and *ACE2* genes and the ratio of expressing cells in the indicated cell types. The color of each bubble represents the level of expression and the size indicates the proportion of expressing cells. **e**, Barplot indicating the percentage of *ACE2* and *TMPRSS2* expressing cells within each cell cluster.

**Fig. 3.**
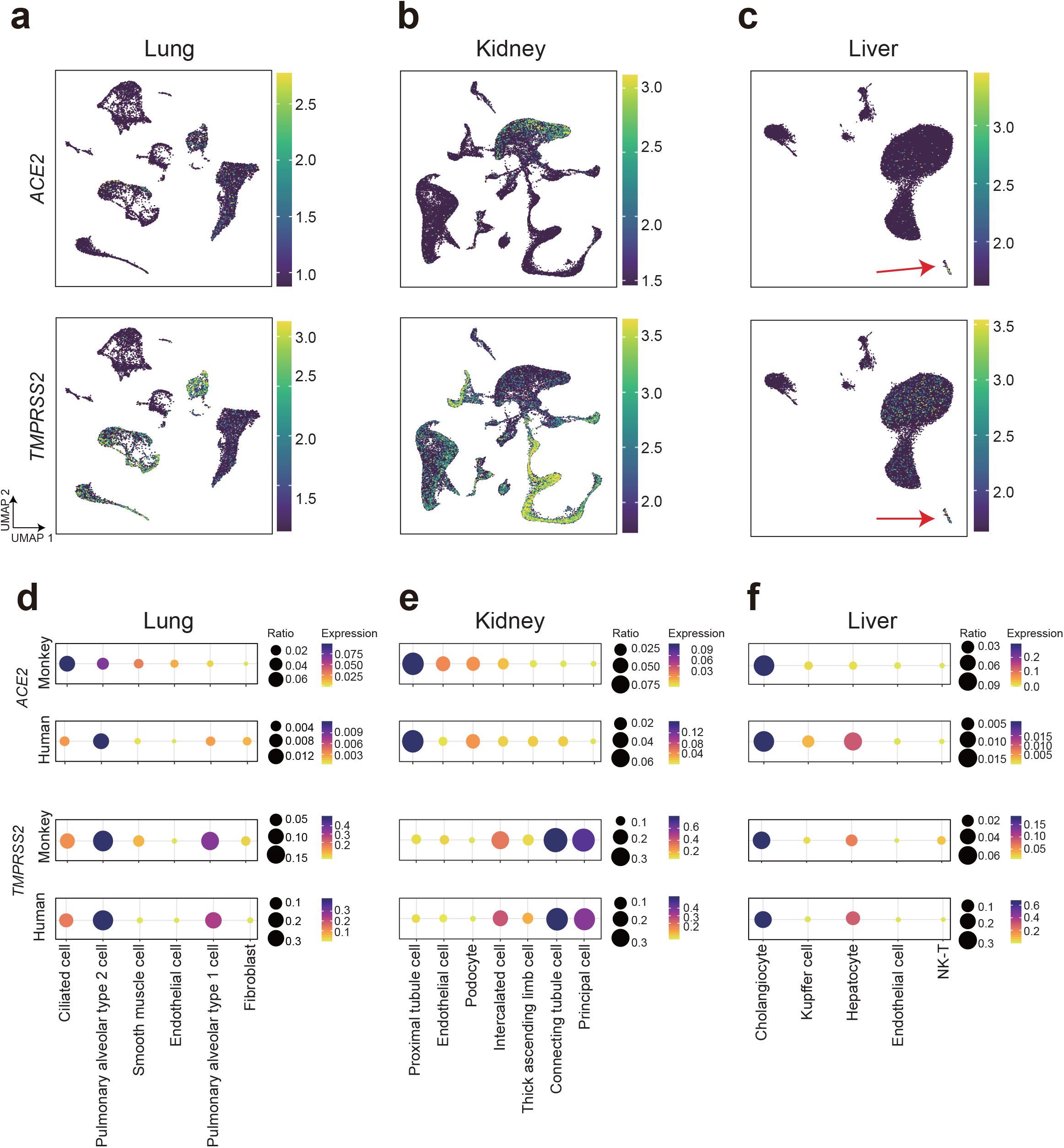
Comparative analysis of *ACE2* and *TMPRSS2* expression between monkey and human. **a-c**, UMAP projection of *ACE2* (top) and *TMPRSS2* (bottom) expression in single cells of monkey lung (**a**), kidney (**b**) and liver tissues (**c**). The red arrow in this panel indicates cholangiocytes. The color of the cells reflects the expression level as indicated in the scale bar. **d-f,** Bubble plots showing the ratio and expression of *ACE2 and TMPRSS2* in the indicated cell types of lung (**d**), kidney (**e**) and liver (**f**) in monkey and human. The color of each bubble represents the level of expression and the size indicates the proportion of expressing cells.

*TMPRSS2* displayed more broadly expressed across cell types in multiple tissues, although it was highest in kidney cells. However, in contrast to *ACE2*, its expression was highest in the distal convoluted tubule, thin limb, intercalated and principal cell 1 and 2 kidney clusters (**Fig. 2b, 2d, 3b lower panel, Extended Data Fig. 3b**). Additionally, significant *TMPRSS2* was observed in both parotid and pancreatic acinar cells, thyroid follicular cells, cholangiocytes and in several lung clusters (**Fig. 2b, 2d, Extended Data Fig. 3**). We then determined which cells co-expressed both genes (*ACE2*^+^/*TMPRSS2*^+^). Notably, the largest overlap between *ACE2* and *TMPRSS2* was observed in the ciliated and club cell clusters of the lung and to a lesser extent the proximal tubule cells of the kidney (**Fig. 2c, 2e**). A smaller overlap was also observed in cholangiocytes and in pancreatic islet cells (**Fig. 2c, 2e**).

Therefore, our data show that *ACE2* and *TMPRSS2* are expressed in a variety of cell types, mainly epithelial cells, within the nine monkey organs/tissues (**Supplementary Table 2a**). The observed heterogeneity of *ACE2* in these cell subtypes also suggests that variations in cell state (e.g. differentiation state, stimulation state or topographical distribution) cause heterogenous expression across an individual tissue. These observations may provide important clues about COVID-19 pathogenesis and symptomatology.

### Comparative analysis of *ACE2* and *TMPRSS2* expression in human and non-human primate

Given the heterogeneous nature of *ACE2* and *TMPRSS2* expression within monkey tissues, we decided to investigate similarities and differences between human and monkey. For this purpose, we retrieved publicly available data from single-cell studies in human (see methods). *TMPRSS2* distribution was similar in cell subtypes of lung, kidney and liver between human and monkey (**Fig. 3d-3f**). However, strikingly, *ACE2* showed distinct patterns among cell subtypes in all three organs between the two species (**Fig. 3d-3f**). The biggest differences were observed in ciliated cells of the lung, which had the highest expression of *ACE2* in monkey, and pulmonary alveolar type 2 cells, which had the highest expression of *ACE2* in human. The function of ciliated cells is to move substances (e.g. cell debris and toxic material) across the surface of the respiratory tract and are commonly targeted by respiratory viruses, whereas pulmonary alveolar type 2 cells have regenerative properties, are crucial for alveolar homeostasis and produce surfactant^24, 25^. In the kidney of both monkey and human, the highest *ACE2* expression was in proximal tubule cells (**Fig. 3e**), which are responsible for electrolyte and nutrient reabsorption. However, renal endothelial cells had higher expression in monkey compared to human. In liver, cholangiocytes had similarly high *ACE2* expression in monkey and human, but hepatocytes showed higher expression and more positive cells in the human (**Fig. 3f**). Considering the heterogenous expression of *ACE2* within the proximal tubule cells in monkey, we revisited the previously analyzed data and were able to sub cluster this population of cells into two (S1 and S3) based on the expression of *SLC5A2* and *SLC7A13*^26^ (**Extended Data Fig. 4, Supplementary Table 2b**). These two genes are sodium and glucose cotransporters involved in glucose reabsorption in the kidney^27, 28^. We did not include thyroid, pancreas or aorta in these analyses because of lack of high-quality available human single-cell datasets. As for the neocortex and PBMC, they have little to no expression of *ACE2* in human (data not shown).

These differences in *ACE2* expression across cell subtypes in the lung, kidney and liver in monkey and human raise the possibility that infection with SARS-CoV-2 in the two species will have different effects.

### *ACE2* correlation analysis across cell types reveals potential therapeutic targets

To shed light on potential mechanisms that could facilitate *ACE2*-mediated SARS-CoV-2 infection, we performed an analysis of the Pearson’s correlation coefficient, based on gene expression in the 44 cell subtypes, to determine what genes are co-regulated with *ACE2* in monkey tissues. Correlated genes were considered as those displaying a coefficient higher than 0.6 with an adjusted *P* value < 0.001. Using these criteria, we observed several genes with marked correlation, including genes that belong to metabolic and developmental pathways and genes involved in the cellular response to xenobiotic stimuli (**Fig. 4a, b**). The highest correlation was observed for transmembrane protein 27 (*TMEM27*, cor = 0.84), a protein involved in trafficking amino acid transporters to the apical brush border of kidney proximal tubules^29^. This is unsurprising considering that *TMEM27* is an important paralog of *ACE2*, and high expression was restricted to kidney cells. DnaJ heat shock protein family (Hsp40) member C12 (*DNAJC12*, cor = 0.78), a gene with a role in immune response processes^30^, had a distribution like *TMEM27*. Importantly, we also observed high correlation with Indoleamine 2,3-dioxygenase 2 (*IDO2*, cor = 0.77), a gene with abundant expression in kidney and liver cells that was also expressed in the lung and other organs. *IDO2* functions during the early phases of immune responses and promotes inflammatory autoimmunity^31, 32^. *ANPEP*, which encodes for alanyl aminopeptidase N, was also co-expressed with *ACE2* in kidney, liver and to a lesser extent in lung too (cor = 0.64), like *IDO2* (**Fig. 4c, d**). Interestingly, *ANPEP* has also been shown to be participate in immune responses, virus receptor activity and in mediating virus entry into host cells^33, 34^.

**Fig. 4.**
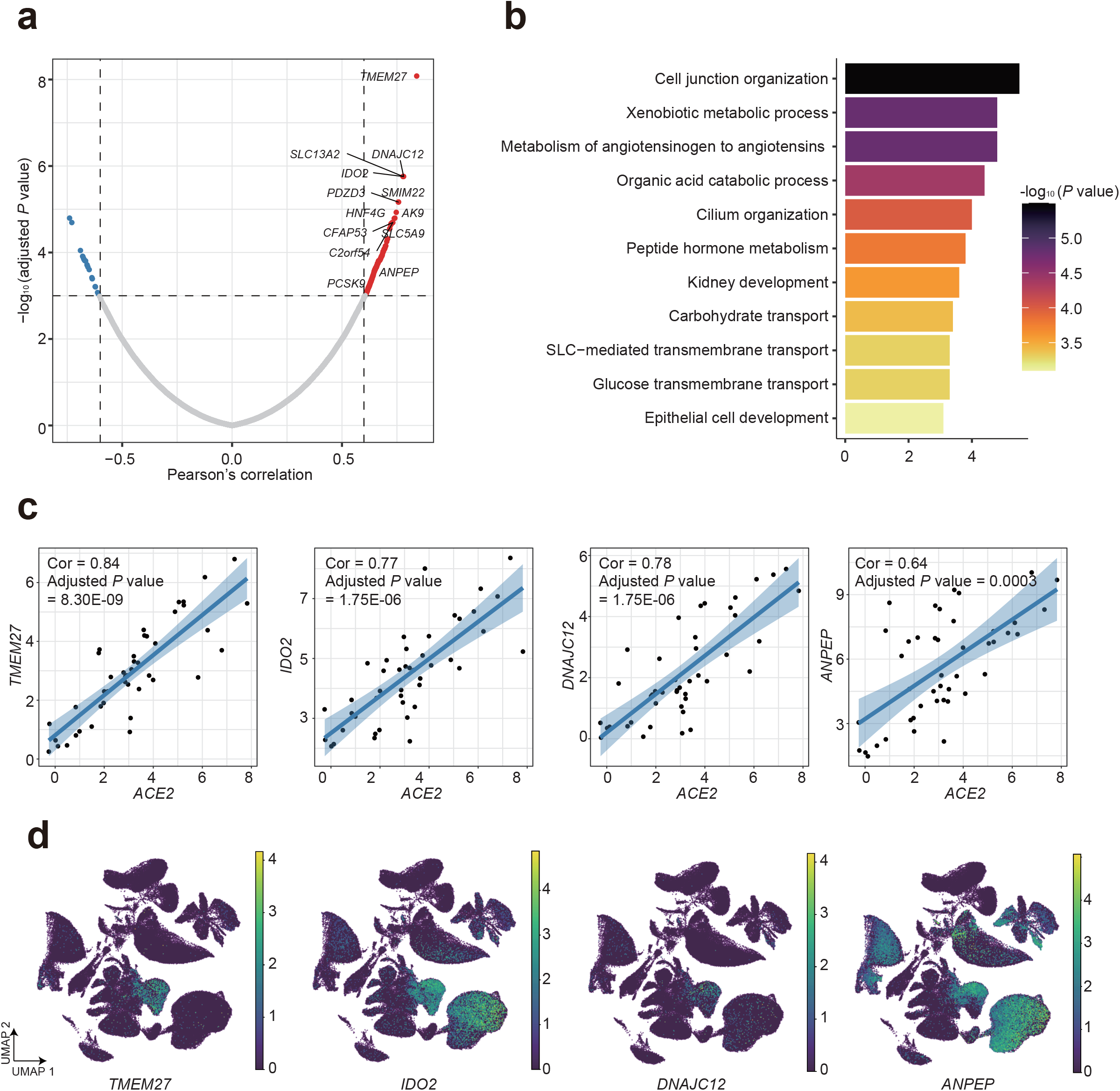
Co-expression analysis of *ACE2* in monkey tissues. **a**, Volcano plot of correlation coefficients (Pearson r^2^) from association tests between *ACE2* and other individual genes. The correlation coefficient for every gene (x-axis) versus the adjusted *P* value (using Benjamini-Hochberg correction; y-axis). The genes indicated in the plot are those with a correlation score > 0.6 and an adjusted *P*-value < 0.001. **b**, Gene ontology analysis of genes that show high expression correlation with *ACE2*. **c**, Scatter plots showing the association between *ACE2* and the indicated genes. The correlation coefficients (Pearson r^2^) and adjusted *P* values are shown in the plots. **d**, UMAP projection of expression of the indicated genes in all single cells.

These data highlight potential therapeutic targets to help in the fight against SARS-CoV-2. Due to their potential co-regulation with ACE2, DNAJC12 and ANPEP it is also possible that they modulate and/or are directly involved in viral entry. Alternatively, depletion of cells expressing *IDO2* and *ANPEP* through a cytopathic effect of the virus could trigger an uncontrolled immune response and contribute to the immune cell exhaustion observed in COVID-19^35^.

### Epigenetic regulation of *ACE2* in each cell subtype of monkey kidney

To understand whether epigenetic mechanisms underlie the heterogeneity of *ACE2* expression in the kidney, as representative for other organs, we employed DNBelab C4 technology to perform high-throughput single-cell ATAC-seq (**Fig. 5a**). After filtering, 6,353 nuclei were used for downstream analysis (**Extended Data Fig. 5a, b, Supplementary Table 4**). We integrated these data with the kidney transcriptomic data described in **Fig. 1** and proceeded to perform Louvain clustering to map all the different cell types within the dataset (**Fig. 5b**). Consistent with the transcriptomic data, our epigenomic mapping identified thick ascending limb cells and proximal tubule cells as the largest kidney clusters (**Extended Data Fig. 2b**). Similarly, smaller clusters of podocytes, principal, intercalated, connected tubule, distal convoluted tubule, thin limb, endothelial and stromal cells were detected (**Fig. 5c, Extended Data Fig. 5c**). Analysis of open chromatin regions revealed discrete peaks in the *ACE2* locus, with the highest signal detected in proximal tubule cells S1 and S3, which are also the highest *ACE2*-expressing cells (**Fig. 5d**). Our approach failed to detect significant signal enrichment in the *ACE2* locus in endothelial cells, possibly related to the low level of expression (**Fig. 5d**). Within the cells of the kidney we observed the highest percentage of *ACE2*^+^ cells in the proximal tubule S3, with a lower percentage in the proximal tubule S1 and endothelial cells (**Fig. 5e**). Motif analysis within the open chromatin regions in *ACE2*^+^ cells demonstrated that these regions were preferentially enriched in STAT1 and 3 and IRF1 binding sites (**Fig. 5f**). These findings suggested that tissue protective cytokines including IL5, IL6, EGF and interferons are acting on these proximal tubule cells S3 to induce *ACE2*. We focused on IL6 because a recent clinical trial has been started that uses anti-IL6 receptor (IL6R) antibodies in the treatment of COVID-19 (http://www.chictr.org.cn/showprojen.aspx?proj=49409). IL6 is a potent regulator of immune responses and can be produced by a variety of interstitial cells including fibroblasts, endothelial cells and more importantly tissue macrophages^36^. Interestingly, we also noticed that distribution of *IL6R* correlated well with *ACE2* in proximal tubule cells (**Fig. 5g, Extended Data Fig. 5d**). In human kidney a similar co-expression pattern was detected (**Fig. 5h**).

**Fig. 5.**
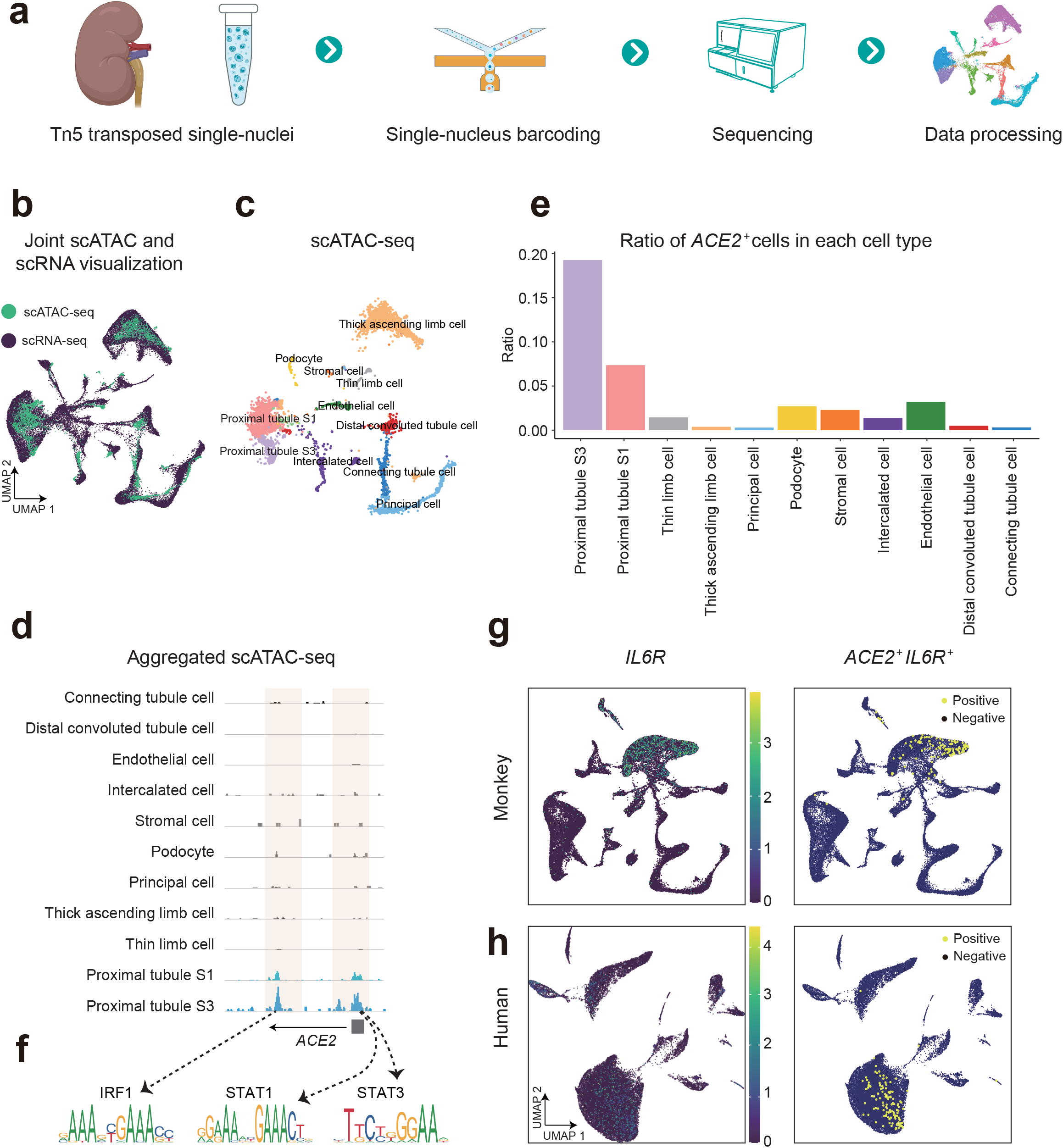
Chromatin accessibility analysis reveals epigenetic regulation of *ACE2* in kidney. **a**, Schematic of experimental design for single-cell ATAC-seq of monkey. **b**, Joint UMAP visualization of kidney single-cell ATAC (scATAC)-seq data with single-cell RNA (scATAC)-seq data. **c**, UMAP visualization of single-cell ATAC-seq data. **d**, IGV visualization of aggregate single-cell ATAC-seq signal in each cell type. **e**, Ratio of *ACE2*^+^ cells in each cell type of kidney. **f**, The transcription factor motifs predicted based on DNA sequence within those regions of the *ACE2* locus. **g-h,** UMAP projection of *IL6R* expression and cells with *IL6R*^+^/*ACE2*^+^ cells in all kidney single cell in monkey (**g**) and human (**h**).

Our observations suggest a potential positive feedback loop between IL6 and *ACE2* expression that can exacerbate COVID-19 disease progression due to increased viral entry and dissemination.

## DISCUSSION

Mammalian tissues and organs are composed of many different cell types that can vary in abundance and cell state. Tissue heterogeneity is only beginning to be unraveled thanks to the advent of single-cell profiling technologies that allow us to precisely map transcriptomic and epigenomic programs. These technologies are revolutionizing our view of human physiology and disease. Great efforts are being made to generate the first version of both human and murine atlases^13, 37^. The mouse is among the most commonly used model organisms in biomedical research but many developmental or pathological aspects are not paralleled in human. Understanding tissue and organ complexity in species that are phylogenetically close to humans is an unmet requirement.

In this study, we have generated a single-cell transcriptomic atlas of nine organs (liver, kidney, lung, pancreas, neocortex, aorta, parotic gland, thyroid and peripheral blood) from cynomolgus monkey. We used this dataset not only to provide fundamental information about the cellular composition of the different tissues tested but also as a platform to dissect the overall expression distribution of the SARS-CoV-2 entry receptor, ACE2, and its serine protease coactivator TMPRSS2^9, 10^. Interestingly, *ACE2* was expressed in multiple epithelial tissues besides the lung, especially the kidney and liver. Other organs of epithelial origin such as the gut have also been implicated in the pathogenesis of the disease^38^. A consequence of this is that the SARS2-CoV-2 virus could infect these organs too, which would explain some of the reported clinical manifestations of COVID-19^2^. By comparing our dataset with publicly available human single-cell RNA-seq data, we have also uncovered significant differences in cell subtypes expressing *ACE2* between human and monkey. We showed different expression patterns for *ACE2* in the lung, where the highest levels were detected in ciliated cells in monkey and pulmonary alveolar type 2 cells in human. Similarly, we observed marked differences in liver, in which monkey hepatocytes displayed higher *ACE2* and a larger number of positive cells compared to the human. We do not know whether these differences will affect the pathogenesis of COVID-19 between these two species. Nevertheless, this is a relevant finding considering that monkeys are a preferred model for studying the effectiveness of drug treatments and of vaccines against the impending COVID-19 pandemic.

Through correlation analysis, we identified new potential mechanisms that could facilitate ACE2-mediated viral infection, including genes previously unreported in the context of SARS-CoV-2 that are involved in stimulating different types of immune responses. We observed high expression of *IDO2* and *ANPEP* in kidney, liver and lung. Expression of these genes can be further induced by viral infection and they have been reported to be immune modulators and/or mediate viral entry^31, 33^. These observations are relevant as they highlight new potential therapeutic vulnerabilities in the current emergency. In this respect, a number of inhibitors of ANPEP are currently being tested in several disease contexts and could serve to prevent the immune cell exhaustion often observed in many severe COVID-19 cases^39^. Similarly, mesenchymal stem/stroma cells (MSC) have immunomodulatory functions that are partly related to IDO2 production ^31^. It is tempting, thus, to speculate that cell therapies based on MSC delivery could ameliorate COVID-19 by normalizing immune function and preventing cytokine storms^40^.

Intriguingly, in our data, we see heterogenous expression of *ACE2* within the individual cell subtypes in six out of the nine monkey organs that we analyzed, which is also the case in the three human organs analyzed. In this regard, we noticed two different cell populations in the kidney proximal tubule, one with higher *ACE2* expression than the other. We performed single-cell ATAC-seq of this organ to understand whether this phenomenon has an epigenetic basis. Analysis of open chromatin regions within the *ACE2* locus revealed the enrichment of STAT1, STAT3 and IRF1 binding sites. These transcription factors have important immune functions and are direct targets of tissue protective and innate immune responses such as Interleukin-6 signaling pathway and interferons. Analysis of *IL6R* distribution showed broad expression within different the *ACE2*^+^ organs in monkey and human. This suggests a link between paracrine IL6 (e.g. secreted by stromal cells including tissue resident macrophages) and enhanced *ACE2* expression across different organs. Higher and more widespread *ACE2* expression could promote increased viral entry. This observation could be very relevant given recent reports describing clinical trials with Tocilizumab, a monoclonal antibody used for IL6R blockade in patients with rheumatoid arthritis^41^, for the treatment of COVID-19 (http://www.chictr.org.cn/showprojen.aspx?proj=49409). IL6 has been related to aging and tissue damage^42^, and this may explain why elderly individuals and those with underlying inflammatory conditions have more severe reactions to SARS-CoV-2 infection (**Fig. 6**). Importantly, high IL6 levels have been detected in plasma from COVID-19 patients^43^. In this context, the proposed enhanced affinity of SARS-CoV-2 for ACE2 compared to SARS-CoV may underlie the differences in the clinical course between the two diseases^44^.

**Fig. 6.**
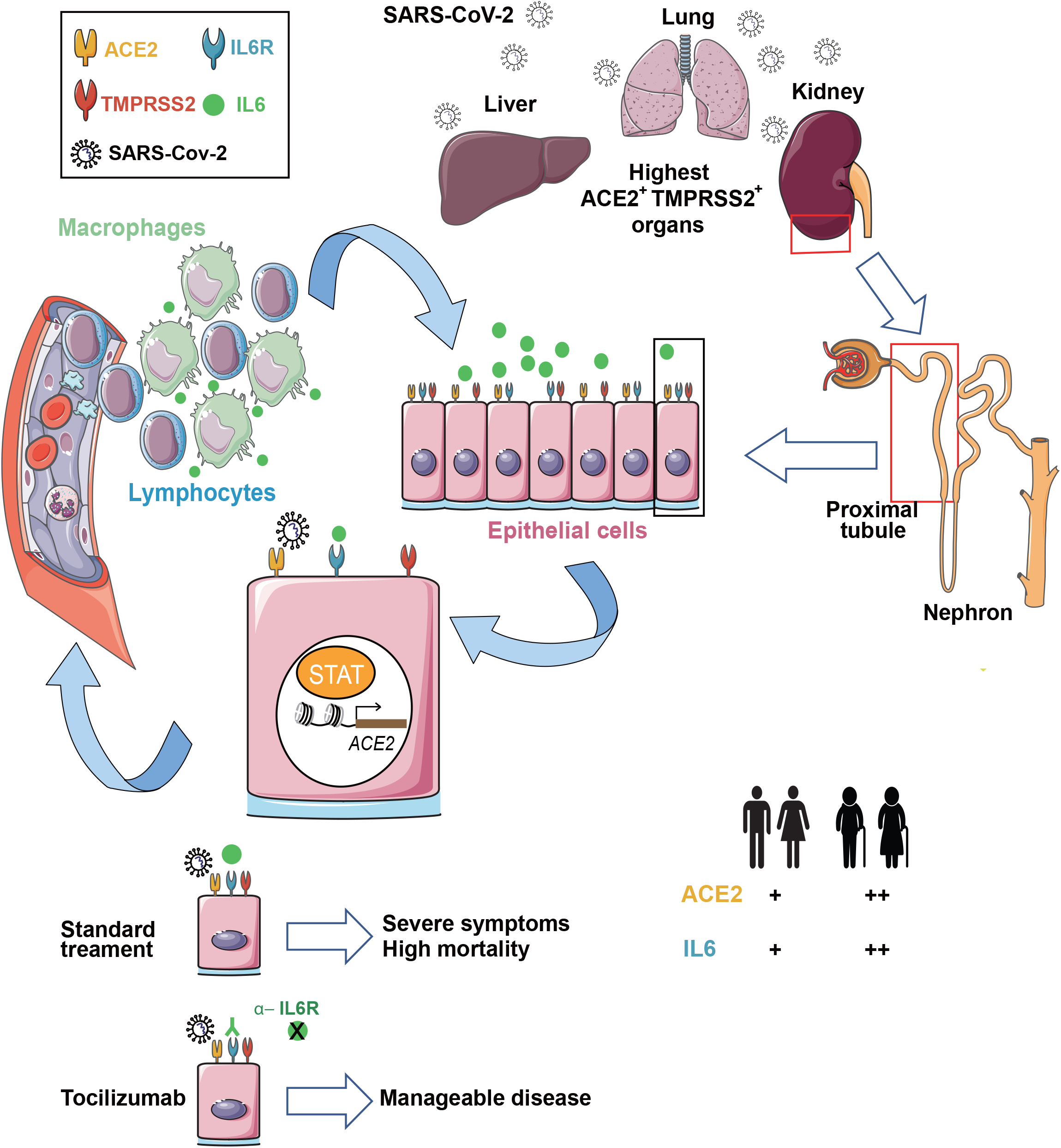
Proposed molecular mechanism for SARS-Cov-2 pathogenesis through reinforced IL6-mediated immune response in monkey and humans. Schematic representation of potential mechanism of SARS-CoV-2 spreading through lung, kidney and liver. Kidney proximal tubule cells within the nephron have the highest expression of ACE2 receptor which facilitates virus entry. After virus contact, IL6R stimulates an immune response that, through the activation of STAT factors, potentiates the paracrine positive feedback loop that facilitates virus spreading. IL6 expression, which is higher in elderly patients and those with inflammatory conditions, is effectively targeted by anti-IL6R monoclonal antibodies leading to a more favourable disease course.

All these observations reveal new potential mechanisms for COVID-19, opening new therapeutic avenues for disease management. However, caution should be exercised when making decisions before additional experimental validation becomes available. Further scrutiny of our datasets may provide new associations useful for understanding COVID-19, and in general will be of utmost relevance for systematic comparisons aiming to understand monkey and human tissue composition and disease vulnerabilities.

## METHODS

### Ethics statement

This study was approved by the Institutional Review Board on Ethics Committee of BGI (permit no. BGI-IRB19125).

### Collection of monkey tissues

A 6-year old female cynomolgus monkey was purchased from Huazhen Laboratory Animal Breeding Centre (Guangzhou, China). The monkey was anesthetized with ketamine hydrochloride (10 mg/kg) and barbiturate (40 mg/kg) administration before being euthanized by exsanguination. Tissues were isolated and placed on the ice-cold board for dissection. Whole organs including lung, kidney, pancreas, liver, brain, thyroid, parotid gland, and aorta were cut into 5-10 pieces, respectively (50-200 mg/piece). Samples were then quickly frozen in liquid nitrogen and stored until nuclear extraction was performed. PBMC were isolated from heparinized venous blood using a Lymphoprep™ medium (STEMCELL Technologies, #07851) according to standard density gradient centrifugation methods. PBMC were resuspended in 90% FBS, 10% DMSO (Sigma Aldrich, #D2650) freezing media and frozen using a Nalgene^®^ Mr. Frosty^®^ Cryo 1°C Freezing Container (Thermo Fisher Scientific, #5100-0001) in a −80°C freezer for 24 hours before being transferred to liquid nitrogen for long-term storage.

### Single-nucleus/cell suspension preparation

We isolated nuclei as previously described^45^. Briefly, tissues were thawed, minced and added to lysis buffer. Lysates were filtered and resuspended in cell resuspension buffer. Frozen PBMC vials were rapidly thawed in a 37°C water bath for ~2 minutes, then quenched with 10 ml 37°C pre-warmed 1X phosphate-buffered saline (PBS, Thermo Fisher Scientific, #10010031) supplemented with 10% FBS. PBMCs were centrifuged at 500 *R.C.F.* for 5 minutes at room temperature. The supernatant was removed, and the cell pellet was resuspended in 3 ml 37°C pre-warmed 1X PBS containing 0.04% bovine serum albumin (BSA, Sangon Biotech, A600903), passed through a 40 μm cell strainer (Falcon, #352340) and then centrifuged at 500 *R.C.F.* for 5 minutes at room temperature. Nuclei or cells were resuspended with cell resuspension buffer at a concentration of 1,000 cells/μl for single-cell library preparation.

### Single-nucleus/cell RNA-seq

The DNBelab C Series Single-Cell Library Prep Set (MGI, #1000021082) was utilized as previously described (Liu et al. 2019). In brief, single-nucleus/cell suspensions were used for droplet generation, emulsion breakage, beads collection, reverse transcription, and cDNA amplification to generate barcoded libraries. Indexed single-cell RNA-seq libraries were constructed according to the manufacturer’s protocol. The sequencing libraries were quantified by Qubit™ ssDNA Assay Kit (Thermo Fisher Scientific, #Q10212). Single-cell ATAC-seq libraries were prepared using DNBelab C Series Single-Cell ATAC Library Prep Set (MGI, #1000021878). DNA nanoballs (DNBs) were loaded into the patterned Nano arrays and sequenced on the ultra-high-throughput DIPSEQ T1 sequencer using the following read length: 30 bp for read 1, inclusive of 10 bp cell barcode 1, 10 bp cell barcode 2 and 10 bp unique molecular identifier (UMI), 100 bp of transcript sequence for read 2, and 10 bp for sample index.

### Single-cell RNA-seq data processing

Raw sequencing reads from DIPSEQ-T1 were filtered and demultiplexed using PISA (version 0.2) (https://github.com/shiquan/PISA). Reads were aligned to Macaca_fascicularis_5.0 genome using STAR (version 2.7.4a)^46^ and sorted by sambamba (version 0.7.0)^47^. Cell versus gene UMI count matrix was generated with PISA.

### Cell clustering and identification of cell types

Clustering analysis of the complete cynomolgus monkey tissue dataset was performed using Scanpy (version 1.4)^48^ in a Python environment. Parameters used in each function were manually curated to portray the optimal clustering of cells. In preprocessing, cells or nuclei were filtered based on the criteria of expressing a minimum of 200 genes and a gene which is expressed by a minimum of 3 cells or nuclei. Filtered data were ln (counts per million (CPM)/100 + 1) transformed. 3000 highly variable genes were selected according to their average expression and dispersion. The number of UMI and the percentage of mitochondrial gene content were regressed out and each gene was scaled by default options. Dimension reduction starts with principal component analysis (PCA), and the number of principal components used for UMAP depends on the importance of embeddings. The Louvain method is then used to detect subgroups of cells. Distinguishing differential genes among clusters were ranked (Benjamini-Hochberg, Wilcoxon rank-sum test). Cell types were manually and iteratively assigned based on overlap of literature, curated and statistically ranked genes. Each tissue dataset was portrayed using the Seurat package (version 3.1.1)^49^ in R environment by default parameters for filtering, data normalization, dimensionality reduction, clustering, and gene differential expression analysis. Finally, we annotated each cell type by extensive literature reading and searching for the specific gene expression pattern.

### Gene correlation and Gene Ontology (GO) term enrichment analysis

The correlation between *ACE2* and other genes was drawn using Pearson correlation coefficient (PCC) with gene expression value merged from cells of the same cell types with the R package psych (version 1.9.12.31). To infer the biological function of highly correlated genes (cor > 0.6 and adjusted P value < 0.001), we performed gene set enrichment analysis using Metascape (https://metascape.org/gp/index.html).

### Differential gene expression analysis

Differential expression analysis between proximal tubule S1 and proximal tubule S3 was performed using the FindMarkers function of the Seurat package (version 3.1.1).

### Single-cell ATAC-seq data processing

Raw sequencing reads from DIPSEQ-T1 were filtered and demultiplexed using PISA (version 0.2) (https://github.com/shiquan/PISA). Peak calling was performed using MACS2 (version 2.1.2)^50^ with options -f BAM -B -q 0.01 –nomodel. The cell versus peak reads count matrix was generated by custom script. To create a gene activity matrix, we extracted gene coordinates for cynomolgus monkey from NCBI, and extended them to ±2 kb region around TSS. The gene activity score matrix was calculated by custom script.

### Single-cell ATAC-seq cell clustering and cell type identification

Cells with low fragments (<1,000) and TSS proportion (<0.1) were removed. Then, filtered data were imported into R and the dimensionality was reduced by latent semantic indexing. Anchors between single-cell ATAC-seq and single-cell RNA-seq datasets were identified and used to transfer cell type labels identified from the single-cell RNA-seq data. We embedded the single-cell ATAC-seq and single-cell RNA-seq data by the TransferData function of Seurat (version 3.1.1).

### Transcription factor motif enrichment analysis

To predict the motif footprint in peaks within the *ACE2* promoter, we extracted genome sequences in the peak region with Seqkit (version 0.7.0)^51^. The sequences were imported into R and were matched with all *Homo sapiens* motifs form JASPAR2018 using matchMotifs function in motifmatchr packages version 1.8.0 with default parameter.

### Human single-cell RNA-seq datasets

All human single-cell RNA-seq data matrix were obtained from publicly available dataset as described: (1) Kidney data from Stewart et al. was download from https://www.kidneycellatlas.org/^52^; (2) Lung data from Madissoon et al. was download from https://www.tissuestabilitycellatlas.org/^53^; (3) Liver data from Aizarani et al. was download from GEO at accession GSE124395 ^54^.

## Code availability

Computer code used for processing the single-cell RNA-seq and single-cell ATAC-seq will be available at https://github.com/brucepan10/NHP-COVID-19.

## Data availability

All raw data have been deposited to CNGB Nucleotide Sequence Archive (accession code: CNP0000986; https://db.cngb.org/cnsa/project/CNP0000986/public/)

## ACKOWLEDGEMENTS

We thank Xiaoyun Huang and Miaomiao Jiang of the Zhiyu Center for Systems Biology, Zhiyu Inc., Shenzhen for their support. This work was supported by National Natural Science Foundation of China (31900466, 31900582), Research and Development Program of China (2018YFA0106903), the Strategic Priority Research Program of the Chinese Academy of Sciences (XDA16030502) and CAS-JSPS Joint Research Projects (GJHZ2093), Natural Science Foundation of Guangdong Province, China (2018A030313379), the Shenzhen National Key Laboratory of Single-Cell Omics (ZDSYS20190902093613831) and Shenzhen Bay Laboratory (SZBL2019062801012). G.V. is supported by Chinese Academy of Sciences President's International Fellowship for Foreign Experts (2020FSB0002). C.W. is supported by Chinese Academy of Sciences President’s International Fellowship Initiative for Postdoctoral Researchers (2019PB0177) and Research Fund for International Young Scientists grant (31950410553). This work is part of the South China Greater Bay Area-Single Cell Consortium under the coordination of M.A.E, L.L., X.X. and Y.H.. This publication is part of the Human Cell Atlas - www.humancellatlas.org/publications/.

## AUTHOR CONTRIBUTIONS

L.L., M.A.E., Y.H. and X.X. conceived the idea. L.L., L.H., Y.L., S.L., X.W. and Y.Yuan. designed the experiment. L.H., Y.L., S.L., X.W., Y.Y., M.C. and C.W.W. collected the tissue samples. C.L., Z.W., Y.Yuan, Y. Yu, M.W., T.W., Y.L., C.W., Y.Z., T.T., Y.H., H.L., L.X., J.X. and M.C. performed the experiments. X.W., T.P., Q.S., L.W., Z.Z., Y.L., S.Z. and S.L. performed the data analysis. L.L., L.H., X.W., C.L., G.V., T.P., C.W. and Y.L. prepared the figures. P.H.M provided critical review of the manuscript. M.A.E., G.V., C.W., Y.L. and L.L. wrote the manuscript with input from all authors. X.X., Y.H., L.L. and M.A.E supervised the entire study. All other authors contributed to the work. All authors read and approved the manuscript for submission.

## EXTENDED DATA FIGURE LEGENDS

**Extended Data Fig. 1.**
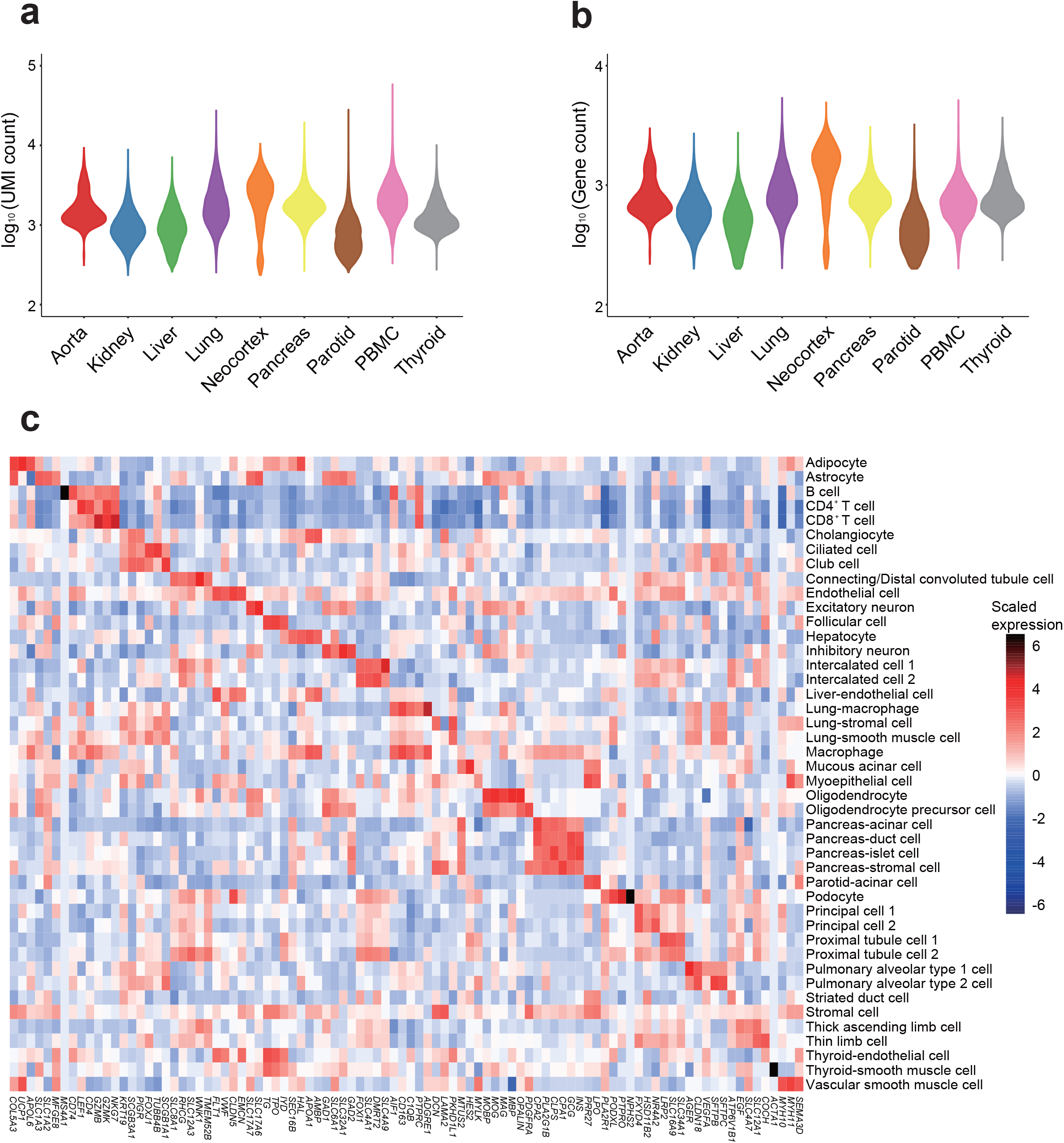
Quality control of the single-cell RNA-seq libraries. **a**, Violin plot showing the number of unique molecular identifiers (UMIs) identified in each tissue. **b**, Violin plot showing the number of genes identified in each organ. (**C**) Heatmap showing the expression of marker genes of the indicated cell type

**Extended Data Fig. 2.**
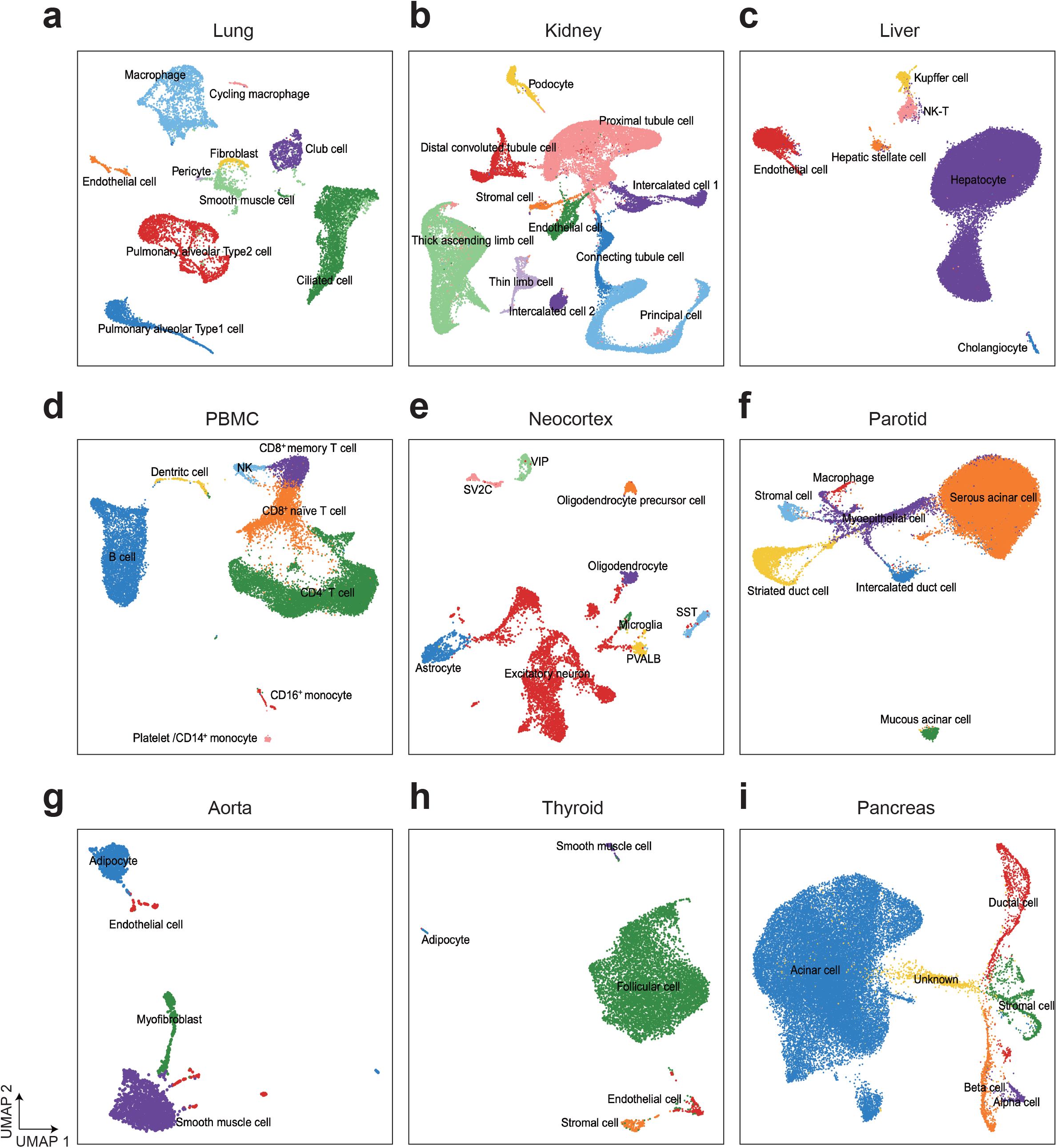
Various cell types identified in each tissue. **a-i**, UMAP visualization of cell clusters in lung (**a**), kidney (**b**), liver (**c**), PBMC (**d**), neocortex (**e**), parotid (**f**), aorta (**g**), thyroid (**h**) and pancreas (**i**). The name of the population corresponding to each cell cluster is indicated in every plot.

**Extended Data Fig. 3.**
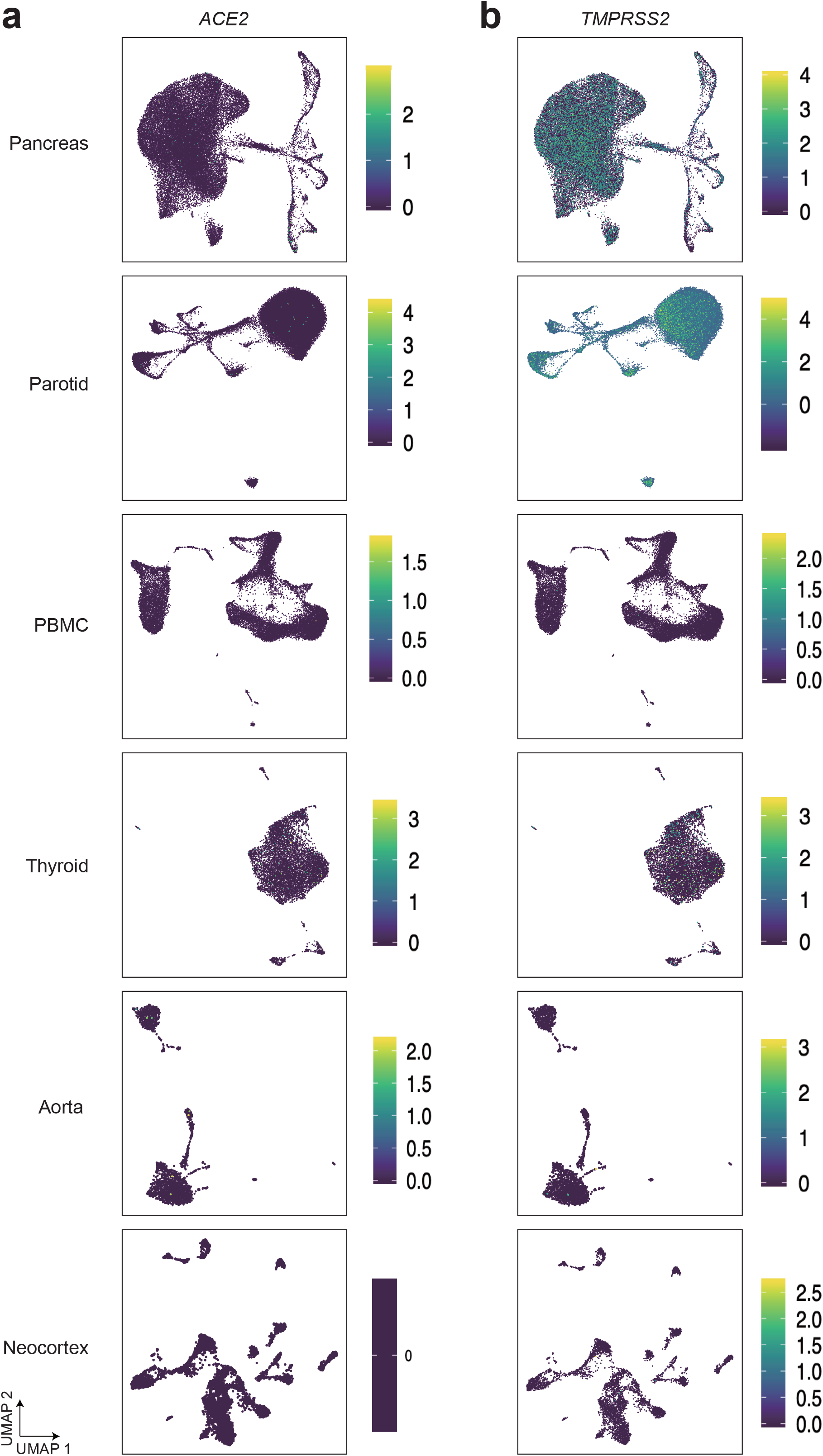
*ACE2* and *TMPRSS2* expression in each tissue. **a-b**, UMAP Projection of *ACE2* (**a**) and *TMPRSS2* (**b**) expression in each tissue.

**Extended Data Fig. 4.**
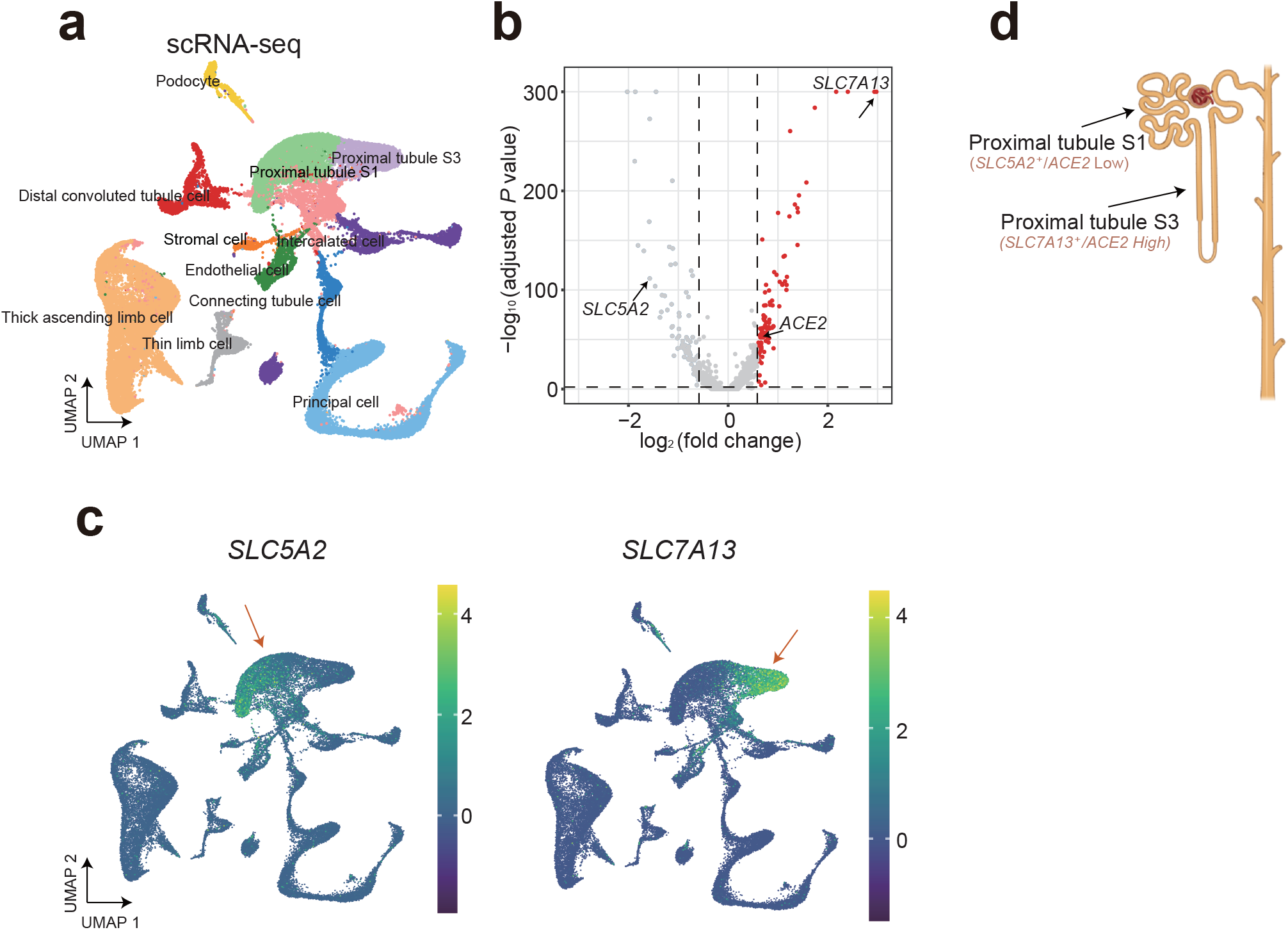
Spatially specific subclusters of proximal tubule cells in monkey kidney. **a**, UMAP visualization of single cells from the kidney tissue, colored by cell types. **b**, Volcano plot showing the differentially expressed genes between proximal tubule S1 and proximal tubule S3 cells. Examples of highly variable genes are indicated. **c**, UMAP projection of expression for the indicated genes in all single cells. **d**, The structure and specific gene expression in kidney tubules. The specific genes and *ACE2* expression level for proximal tubule S1 and proximal tubule S3 cells are indicated.

**Extended Data Fig. 5.**
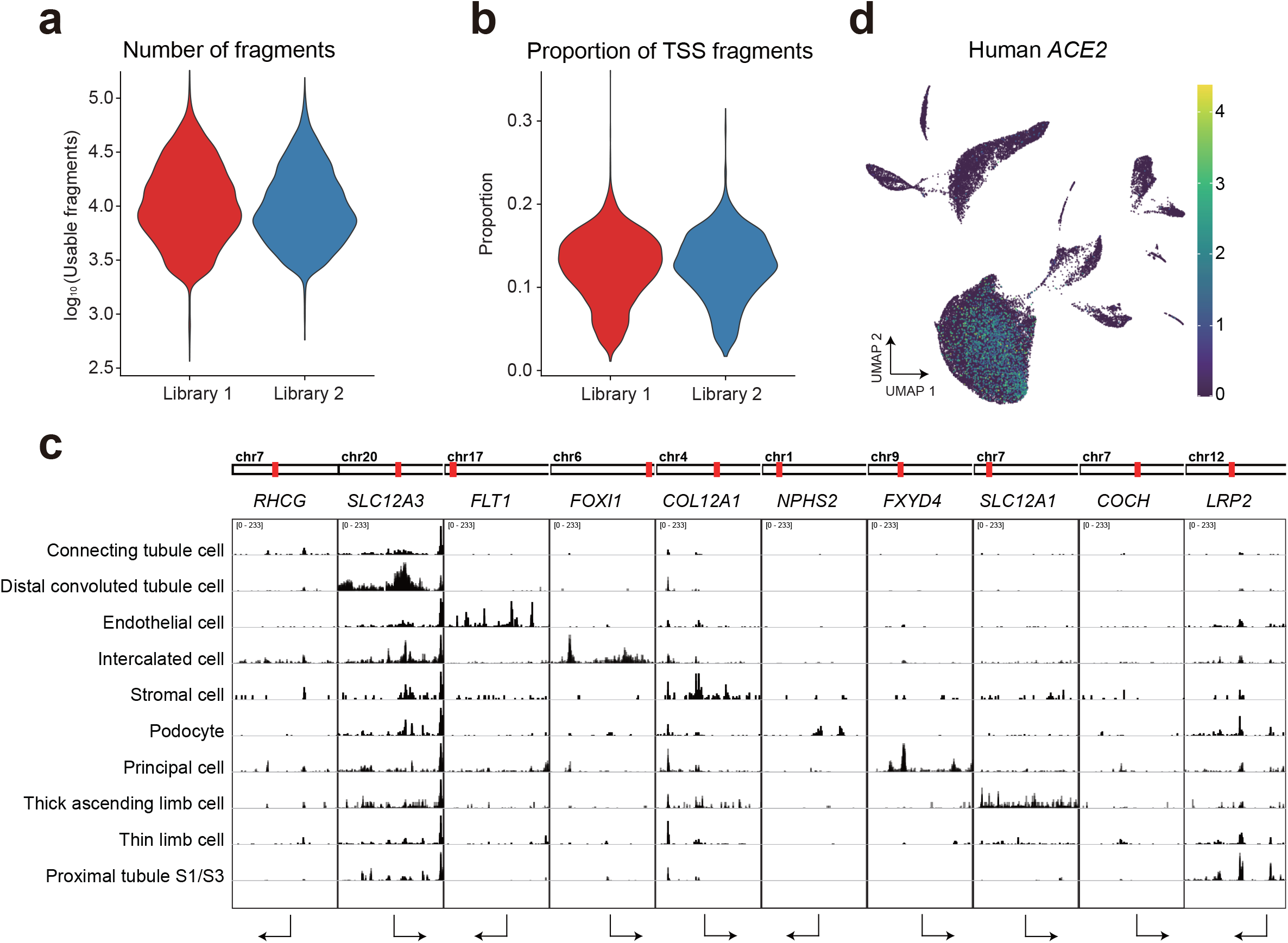
Quality control of single-cell ATAC-seq data. **a**, Number of fragments captured in all cells of the two single-cell ATAC-seq libraries. **b**, Proportion of TSS fragments in all cells of the two single-cell ATAC-seq libraries. **c**, IGV visualization of specific accessible chromatin in each cell type. **d**, UMAP projection of *ACE2* expression in human kidney.

